# 1 Hz rTMS Is Not Inherently Inhibitory: Current Direction Determines Aftereffects and Reliability

**DOI:** 10.64898/2026.06.29.732840

**Authors:** Carolina Kanig, Mirja Osnabruegge, Leo Tomasevic, Berthold Langguth, Wolfgang Mack, Stefan Schoisswohl

**Affiliations:** Department of Psychiatry and Psychotherapy, University of Regensburg, Universitätsstraße 84, 93053 Regensburg, Germany; Department of Human Sciences, University of the Bundeswehr Munich, Werner-Heisenberg-Weg 39, 85577 Neubiberg, Germany

**Keywords:** TMS, rTMS, MEP, cortico-spinal excitability, current direction, reliability, plasticity

## Abstract

**Objective:** Aftereffects of 1 Hz repetitive transcranial magnetic stimulation (rTMS) often differ within and between subjects and thus show low reliability. In this study we investigated the mean and individual aftereffects of 1 Hz rTMS using two opposing current directions, their reliability and potential influences of current direction, participants’ sex and state on cortical excitability modulations.

**Methods:** Thirteen healthy, right-handed participants underwent four experimental sessions separated by at least 7 days receiving 2000 pulses of suprathreshold 1 Hz rTMS over the primary motor cortex per session. Two sessions were conducted with an induced current direction of anterior-posterior – posterior-anterior (AP–PA) and two sessions with a PA-AP current direction. Before and after rTMS, 100 single TMS pulses were administered with the respective current direction and electromyography was recorded from the first dorsal interosseous. Questionnaires on demographic data and subjective ratings were completed during the experiment.

**Results:** Linear mixed effect model analysis revealed that 1 Hz rTMS induced an excitatory aftereffect when applied with the PA-AP current direction, and no aftereffect with AP-PA. There was a substantial interindividual variability with only three subjects showing an inhibition to 1 Hz rTMS overall. Also, current direction was the only predictor of rTMS aftereffect. Reliability values of these aftereffects were in the poor to moderate range.

**Conclusions:** Current direction plays a crucial role in determining 1 Hz rTMS aftereffects. Reliability was found to be moderate at best. Additional to current direction, more factors need to be considered to tailor the 1 Hz rTMS aftereffects individually.

## Introduction

Repetitive transcranial magnetic stimulation (rTMS) is a non-invasive neuromodulation method capable to alter cortical and cortico-spinal excitability (Pascual-Leone et al., 1994; Wassermann et al., 1996). Even single rTMS sessions can induce transient excitability changes that persist beyond stimulation offset, reflecting short-term neuroplasticity mechanisms (Touge et al., 2001). Typically, modulations of cortico-spinal excitability from pre to post rTMS are measured via the application of single TMS pulses over the primary motor cortex (M1) while recording electromyography (EMG) responses of the corresponding contralateral hand muscle (Klomjai et al., 2015). With suprathreshold TMS pulses, the hand muscle can contract, resulting in a motor evoked potential (MEP) (Di Lazzaro et al., 2004; Spampinato et al., 2023). The implementation of rTMS protocols in experimental and clinical research (Chen et al., 1997; Di Lazzaro et al., 2008; Schoisswohl et al., 2021), as well as treatment of neuropsychiatric diseases (Lefaucheur et al., 2020) is widespread. However, the induced aftereffects have not consistently proven to be stable and their reliability remains limited (Kanig et al., 2023; Prei et al., 2023). Especially protocols that are assumed to exhibit long term depression (LTD) -like mechanisms and are intended to inhibit cortical and cortico-spinal excitability, like 1 Hz rTMS (Chen et al., 1997), sometimes fail their purpose (Bäumer et al., 2003; Kanig et al., 2025; Modugno et al., 2003; Weisz et al., 2012). Hereby, responder rates with MEP amplitude as outcome parameter vary vastly (Goetz et al., 2016; Prei et al., 2023) up to 75 % (Wassermann et al., 1996).

It is known that application-related and subject-related factors can not only reduce the desired aftereffect, but also lead to opposite rTMS aftereffects (Guerra et al., 2020; Ridding & Ziemann, 2010). But their full extent is yet to investigate. However, individual variability highlights the limitations of applying the 1 Hz rTMS protocol on a large scale, especially in a therapeutic context, which is why we wanted to investigate it further. Within subjects, most prominently the cognitive state has been found to influence responses to TMS and rTMS (Sack et al., 2024). Between subjects, the factor sex for example seems to influence variability (Pitcher et al., 2003). Among application-related factors, particular attention has been given to the induced current direction, since differences in outcomes based on the applied current direction in both TMS (Casula et al., 2018; Davila-Pérez et al., 2018; Kammer et al., 2001; Osnabruegge et al., 2026) and rTMS (Goetz et al., 2016; Kanig et al., 2025; Schoisswohl et al., 2023; Sommer et al., 2013) have been observed. Yet, for biphasic pulses that are in standard use for 1 Hz rTMS, only two publications investigated differences in cortico-spinal aftereffects (Kanig et al., 2025; Sommer et al., 2013).

In both basic science and clinical studies, 1 Hz rTMS is frequently used to alter cortical plasticity according to its predicted outcome (Lefaucheur et al., 2020; Patel et al., 2020). But since many influential factors can hamper the predicted aftereffects (Guerra et al., 2020), with this study we not only wanted to show the influence of current direction on 1 Hz rTMS aftereffects, but also show their reliability and exploratively analyse the individual responses. Additionally, we wanted to analyse which influence individual demographical and cognitive state parameters such as current direction, the subjects’ sex, attention, excitement or tiredness have on the observed aftereffects in order to better understand sources of variability and identify factors that may contribute to more reliable stimulation effects.

## Methods

The study protocol was reviewed and approved by the local ethics committee of the University of Regensburg (ethical approval number: 21–2662–1–101). All participants gave written informed consent prior to study start, which was conducted in compliance with the ethical principles of the Declaration of Helsinki.

### Participants and exclusion criteria

The sample comprised of 15 healthy, right-handed (Edinburgh Handedness Inventory: *M* = 83.41, *SD* = 15.59; Oldfield, 1971), German-speaking adults (9 female; 20–42 years, *M* = 25, SD = 6). The mean intelligence quotient was 110.33 (*SD* = 11.71) as assessed by the German version of the multiple-choice vocabulary intelligence test, MWTB-Q (Lehrl, 2018), and no participant had neurological, psychiatric, or severe somatic disorders as confirmed by clinical assessment. Exclusion criteria included depressive symptoms as assessed by the Major Depression Inventory, MDI (Bech et al., 2001), stimulation intensity > 80 % maximum stimulator output, psychoactive substance use, and contraindications for TMS (e.g., ferromagnetic implants, severe brain injury, epilepsy). Participants received monetary compensation. In two of the 15 subjects, EMG data was very noisy with no MEPs identifiable (mean amplitude to baseline ratio in one condition respectively was 1.05 and 1.07). Therefore, these two subjects were removed from all analyses.

### Procedure

The study comprised a pre-session and four experimental sessions. During the pre-session, eligibility criteria were verified, written informed consent was obtained, and questionnaires such as the International Physical Activity Questionnaire; IPAQ (Hagströmer et al., 2006), MDI (Bech et al., 2001), MWTB-Q (Lehrl, 2018), and on overall health were completed. All stimulations in the experimental sessions were conducted with either inducing an anterior-posterior – posterior-anterior (AP–PA) or posterior-anterior – anterior-posterior (PA–AP) current direction in the brain; the starting condition was randomized a priori (6 participants AP–PA, 7 PA–AP), and participants were blinded to the current direction throughout the study. Sessions 1 and 2 were performed with the same current direction, whereas sessions 3 and 4 were conducted with the opposite direction, while all were scheduled at least one week apart and at the same time of day.

At the beginning and end of each session, participants rated subjective attention, excitement, and tiredness on visual analogue scales (1–10). Before stimulation, subjects were instructed to remove electronic devices, sit relaxed and still, and fixate a cross during stimulation. EMG was recorded continuously from the right first dorsal interosseous (FDI), abductor pollicis brevis (APB) and abductor digiti minimi (ADM) muscles.

TMS was delivered over left primary motor cortex (M1) using a neuronavigated, cobot-assisted setup. The individual motor hotspot for the FDI was identified with the AP-PA current direction using a grid-based semi-automated procedure (see Agboada et al., 2023), targeting high and stable FDI MEPs with minimal APB and ADM activity at the lowest effective intensity. This motor hotspot was used for all sessions and both current directions to ensure comparable coil positioning and electric field orientation. Resting motor threshold (RMT) was defined as the minimum intensity eliciting MEPs ≥ 50 µV in 50 % of trials (Rossini et al., 2015).

Each session included 100 single TMS pulses at 110 % RMT (ISI 10 ± 2 s) before and after 1 Hz rTMS (2000 pulses, 110 % RMT) delivered over the individual motor hotspot. Stimulation intensity and duration were chosen to maximize rTMS aftereffects (Fitzgerald, 2002; Lang et al., 2006; Peinemann et al., 2004). All pulses were masked with individualized noise (70 % white noise, 30 % TMS click, ≤ 85 dB SPL) created with the TMS Adaptable Auditory Control (TAAC) toolbox (Russo et al., 2022) in Matlab (Matlab R2018b; Mathworks, USA), and delivered through ER-3C 10 Ω Insert Earphones (Etymotic Research Inc., USA) together with an iPod Touch 7th Generation (Apple Inc., Cupertino, California, USA). After each session, side effects were surveyed via a dedicated questionnaire.

### Electromyography (EMG)

We recorded activity of the FDI, APB and ADM of the right hand with Ag/AgCl surface electrodes in bipolar belly-tendon montage (GND: processus styloideus ulnaris) in combination with ActiCHamp with BIP2AUX adapters (Brain Products GmbH, Germany). Raw EMG signal from the Brain Vision Recorder (Brain Products GmbH, Germany) was read into and preprocessed with Matlab (R2021b; Mathworks, USA) and filtered with 4th order two-way butterworth filter with high-pass 10 Hz and low-pass 500 Hz. EMG peak-to-peak amplitude of the FDI was extracted between 20 and 50 ms after the TMS pulse per subject, current direction and day for the 100 TMS trials pre and post rTMS. Trials with high EMG activity, exceeding ± 100 µV, indicating a contracted muscle, 100 to 1 ms before the pulse were excluded.

### Transcranial magnetic stimulation (TMS)

Magnetic stimulation was delivered using a MagPro X100 with MagOption and a figure-of-eight Cool-B65 A/P coil (MagVenture A/S, Farum, Denmark). Biphasic pulses (∼280 µs) were applied to ensure consistent conditions between rTMS and single-pulse measurements (Peinemann et al., 2004). The coil was positioned over the hotspot 45° to the midline using neuronavigation (Localite GmbH, Bonn, Germany) and a collaborative robotic coil holder (Axilum Robotics, Schiltigheim, France), maintaining placement within 2 mm across sessions (Agboada et al., 2023). Two current directions were tested: the *default* setting, inducing an AP–PA current flow in the brain, and the *reversed* setting of the stimulator, inducing a PA–AP current flow in the brain.

### Statistical analyzes

EMG and demographic data were analyzed in R (R Core Team, Austria, Version 4.3) with the packages lme4 (Bates et al., 2015), lmerTest (Kuznetsova et al., 2017), emmeans (Russell V. Lenth, 2023), irr (Gamer et al., 2019) and ordinal (Christensen, 2023). Plots were generated using ggplot2 (Gómez-Rubio, 2017). We analyzed MEP amplitudes with linear mixed effect models. In a stepwise selection approach, we compared models with the predictors *time* (pre vs post 1 Hz rTMS), *day* (1 vs 2), *current direction* (AP–PA vs PA–AP) and their interaction against the intercept model without predictors. Model fit was evaluated and a more complex model was only retained when AIC and BIC decreased and the Likelihood Ratio test reached significance level (Hastie, 2017). Marginal and conditional *R*^2^ were calculated to estimate explained variance (Nakagawa et al., 2017). The factor subject was implemented as random intercept in all models, so that the full model was as follows: *data ∼ time + day + current direction + time*day*current direction + (1*| *subject)*. Fixed effects were evaluated via expected mean squares approach and concurrent contrasts with post-hoc Tukey t-tests and adjusted p-values. Two-way mixed effects intraclass correlation coefficients (ICCs) with absolute agreement (Koo & Li, 2016) were computed on the mean MEP amplitude per subject and condition, overall and separately per current direction.

To exploratory investigate the impact of demographic and individual variables on the 1 Hz rTMS aftereffect, we computed an ordinal logistic regression with the model: *response ∼ current direction + sex + attention (pre) + excitement (pre) + tiredness (pre)*. Hereby, the dependent variable was defined as 1 Hz rTMS aftereffect direction response: inhibition, no change or excitation. It was calculated via kmeans from the stats package for each subject and condition (day 1 and 2 for current direction AP-PA and PA-AP) on the pre and post (time) data. The variables were used to provide an overview of individual (demographics) and experimental (current direction) influencing factors that are discussed in the current literature (Guerra et al., 2020; Ridding & Ziemann, 2010). We first tested whether predictors had zero and near zero variances with the caret package (Kuhn, 2008) as well as collinearity between predictors with the performance package (Lüdecke et al., 2021). Subsequently, we investigated significance of the predictors from the adjusted model via likelihood ratio test (car package; (Fox & Weisberg, 2019)) and their odds ratios. For all analyzes, the level of significance was set at α = .05.

## Results

### rTMS aftereffects

In analysis of EMG amplitude, the fixed effects of *time, current direction* and *day* were significant using the expected mean square approach. Also the interaction effect of *time* and *current direction* reached significance. Marginal *R*^2^ was 0.0167 and conditional *R*^2^ 0.0851. Post-hoc Tukey t-tests revealed that EMG amplitude was higher after 1 Hz rTMS (M = 507, SE = 56) than before (M = 414, SE = 56, t(9335) = 6.464, p < 0.0001), higher when applied with the PA-AP current direction (M = 522, SE = 56) than the AP-PA current direction (M = 400, SE = 56, t(9340)= -8.386, p < 0.0001) and higher on the first day of stimulation (M = 487, SE = 56) than at the second day (M = 434, SE = 56, t(9336)= 3.641, p = 0.0003). The interaction of *time* and *current direction* was mainly driven by the increase of EMG amplitude following 1 Hz rTMS using PA-AP current direction: The EMG amplitude after rTMS was higher (M = 616, SE = 56.9) than before rTMS in the PA-AP current direction condition (M = 428, SE = 56.9, t(9335) = 9.306, p < 0.0001), higher than before AP-PA rTMS (M = 400, SE = 56.9, t(9336) = -10.549, p < 0.0001) as well as higher than after AP-PA rTMS (M = 399, SE = 57, t(9339) = -10.472, p < 0.0001). **Figure 1** depicts the estimated means and the respective confidence intervals.

**Figure 1.**
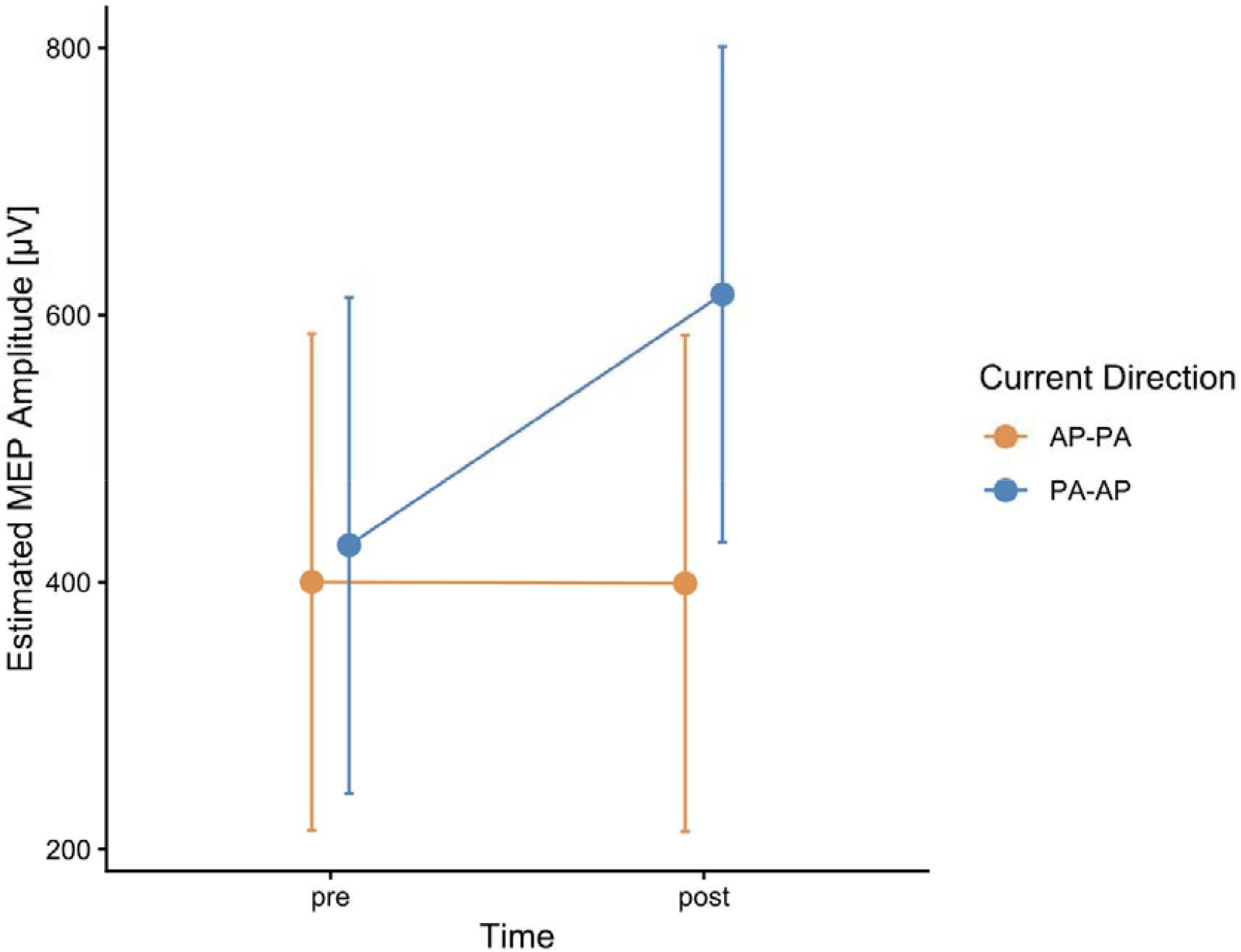
Interaction effect of time and current direction. *Notes*. The interaction effect resulting from the linear mixed effect model is depicted via estimated means of the MEP amplitude (dots) and their 95 % confidence intervals (whiskers). The orange line depicts values from the induced AP-PA current direction and the blue line the PA-AP current direction.

Kmeans clustering revealed that from 52 data points (13 subjects and 4 conditions each) 39 were categorized into “no change”, 8 into “excitation” and 5 into “inhibition”, which is depicted in **Figure 2**. Only three of the thirteen subjects were found to exhibit an inhibition aftereffect response to 1 Hz rTMS at all. With induced AP-PA current direction, four data points were found to be inhibitory and with PA-AP current direction only one in one subject.

**Figure 2.**
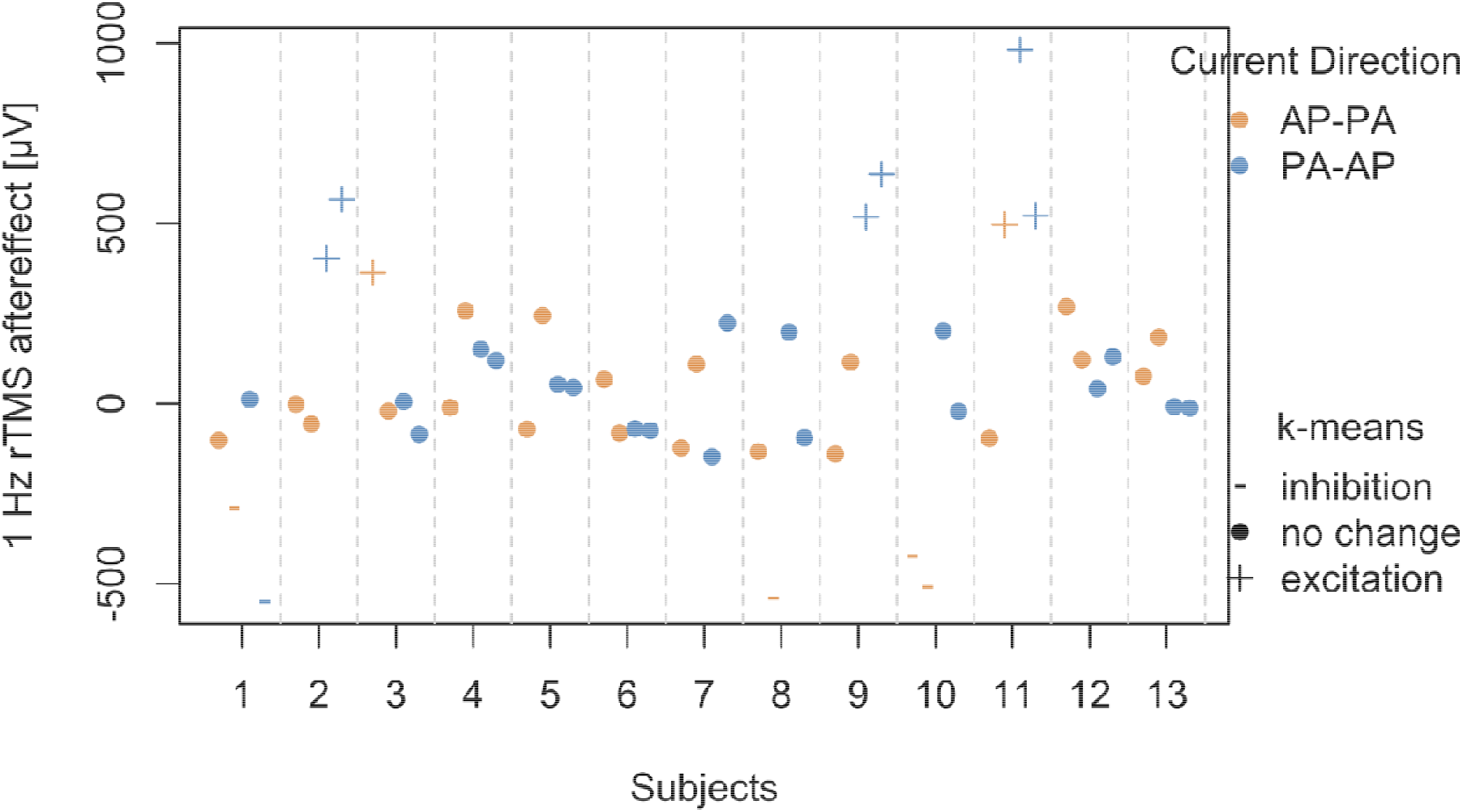
Cluster identified by k-means. *Notes*. Mean 1 Hz rTMS aftereffects per subject and current direction were divided into three clusters via k-means clustering. Minus signs represent an inhibitory aftereffect, dots show no change and plus signs indicate an excitatory 1 Hz rTMS aftereffect. Aftereffects of the sessions with induced AP-PA current direction are depicted in orange and for the PA-AP current direction in blue.

### rTMS reliability

ICC analyses of mean rTMS aftereffect MEP amplitude revealed an overall reliability of rTMS of 0.561 (95 % CI [0.225; 0.777], p < 0.01). When splitting up for rTMS per current direction, the aftereffect of 1 Hz rTMS with AP-PA current direction had an ICC of 0.342 (95 % CI [-0.27; 0.745], p = 0.127) and the aftereffect of 1 Hz rTMS with PA-AP current direction resulted in an ICC of 0.664 (95 % CI [0.222; 0.883], p < 0.01). The individual stability is shown via combined violin and box plots with lines between the individual values in **Figure 3. Figure 4** depicts the mean aftereffects of each subject with their respective differences per current direction in Bland-Altman-Plots (Martin Bland & Altman, 1986).

**Figure 3.**
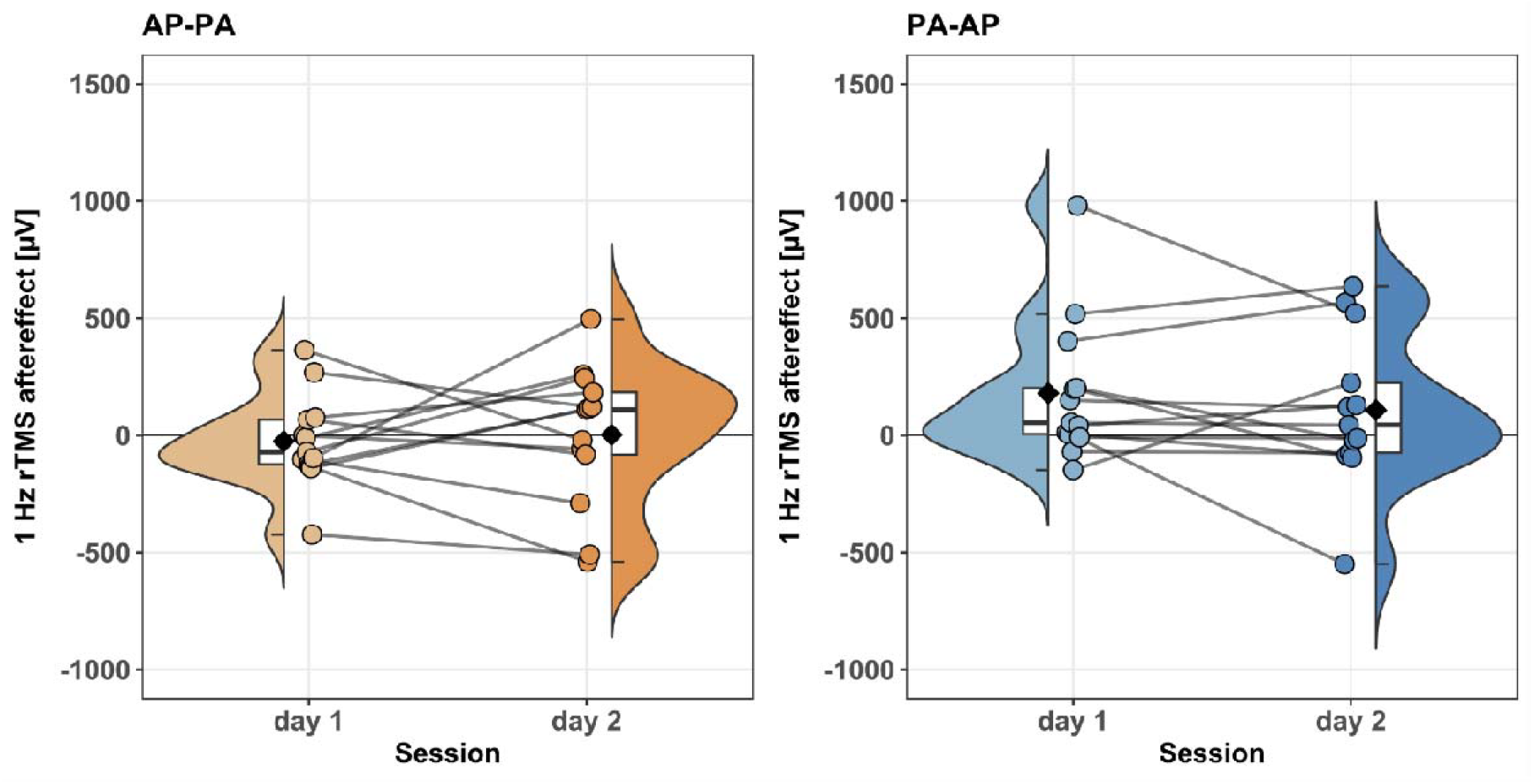
Violin and box plots with individual data points and combining lines per current direction. *Notes*. The left plot depicts the 1 Hz rTMS aftereffects in orange color for the induced AP-PA current direction per subject (dots) on the first and second session, separated by seven days. Additionally, the box plot and violin plot of this sample are depicted as halves corresponding to the individual data. Lines between the dots connect the respective subjects’ data. Black diamonds represent the mean values of the sample. The right plot depicts the corresponding data for the induced PA-AP current direction 1 Hz rTMS sessions in blue color.

**Figure 4.**
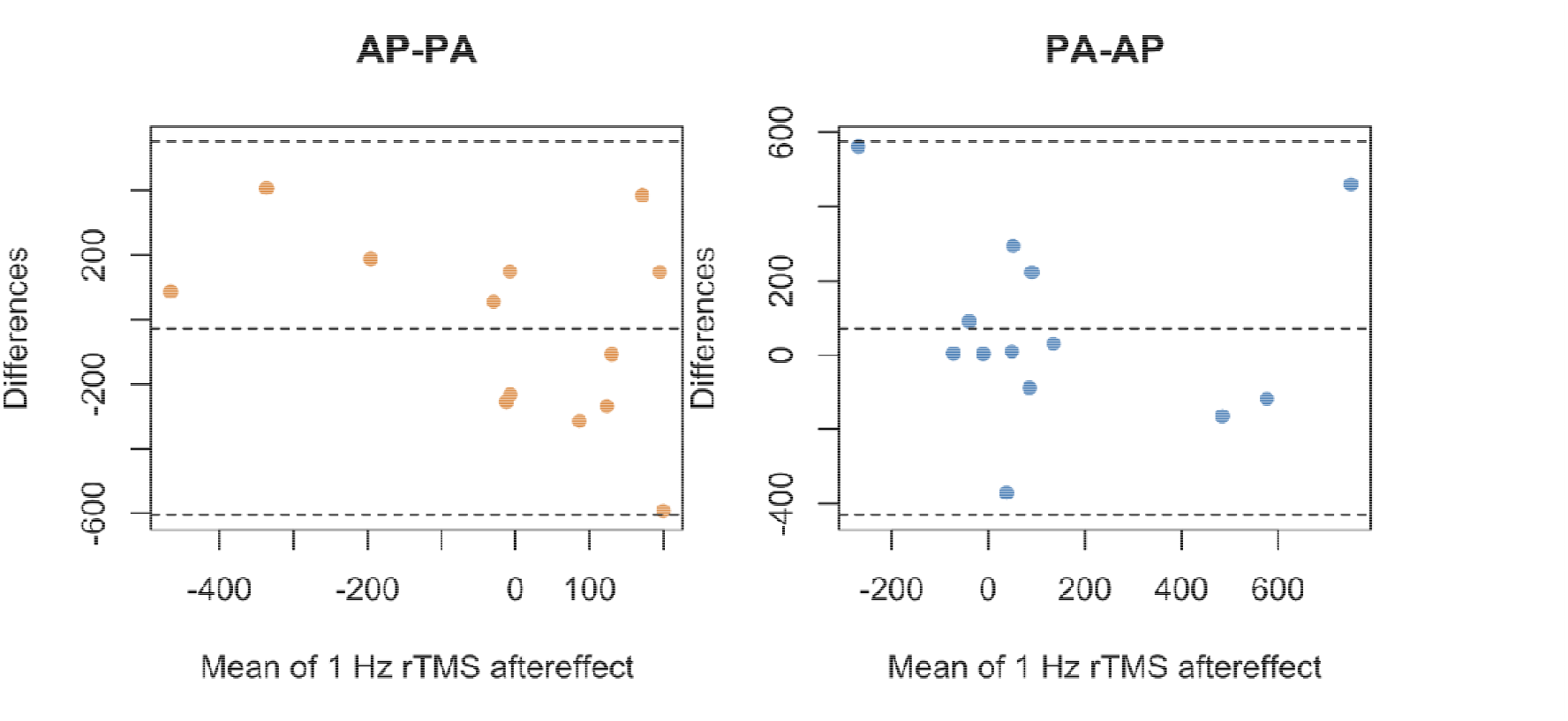
Bland-Altman-Plots per current direction. *Notes*. The mean 1 Hz rTMS aftereffects resulting from the induced AP-PA (left plot, orange dots) and PA-AP (right plot, blue dots) current direction per subject are plotted against their differences from the overall mean 1 Hz rTMS aftereffect per current direction. The dashed lines in the middle depict the mean difference and the outermost dashed lines refer to the 2.5 % and 97.5 % percentile point of the standard normal distribution (± 1.96 SD).

### rTMS influences

None of the predictors from the original investigated ordinal logistic regression model *response ∼ current direction + sex + attention (pre) + excitement (pre) + tiredness (pre)* showed variances that were near to zero (frequency ratios [1 – 1.6154], percent of unique data [3.8462 – 19.2308]). Multicollinearity analyses revealed adjusted variance inflation factors (VIF) below 4 for all the variables.

Type II likelihood ratio tests showed an influential effect of *current direction* (χ^2^(1) = 4.5461, p < 0.05) on the 1 Hz rTMS aftereffect response. The odds ratio of the estimate revealed that the induced PA-AP current direction increased the probability for an excitation response (OR = 5.34, 95 % CI [1.26; 29.7], p < 0.05). All other factors failed to reach significance.

## Discussion

In a within-subject-design study with 2000 pulses of 1 Hz rTMS applied with the induced current directions AP-PA and PA-AP, twice respectively and in randomized order, we investigated the mean and individual aftereffects as well as reliability of the low-frequency rTMS protocol. We found excitatory aftereffects on group level for the PA-AP current direction with low to moderate reliability. Only three subjects exhibited an inhibitory aftereffect after 1 Hz rTMS, with four out of five cases with the induced AP-PA current direction. Additionally, we examined whether current direction, the subjects’ sex, attention, excitement or tiredness influenced these aftereffects and found current direction as the sole predictor.

1 Hz rTMS aftereffects in this experiment showed that the commonly presumed inhibitory protocol exhibited excitatory effects at group-level in the condition with induced PA-AP current direction. The AP-PA current direction did not evoke changes in MEP amplitudes. Although many 1 Hz rTMS applications show inhibitory responses (Fitzgerald, 2002; Goetz et al., 2016; Maeda et al., 2000; Romero et al., 2002), some fail to do so (Daskalakis et al., 2006; Modugno et al., 2003). It may be that our results showed an excitatory effect, because contrary to other 1 Hz rTMS applications we stimulated the healthy primary motor cortex with suprathreshold instead of subthreshold stimuli as (Maeda et al., 2000), (Romero et al., 2002) and (Goetz et al., 2016) did, although higher reduction of MEP amplitude was observed for higher intensity probing pulses (Goetz et al., 2016). Yet, suprathreshold intensities (115 % RMT) of 1 Hz rTMS were found to inhibit cortico-spinal excitability more than subthreshold (85 % RMT) intensities (Fitzgerald, 2002). Also, we applied a higher number of pulses during rTMS than previous literature (Fitzgerald, 2002; Goetz et al., 2016; Maeda et al., 2000; Romero et al., 2002). Although (Maeda et al., 2000) reported higher inhibition aftereffects for the higher number of pulses, since these effects on cortico-spinal excitability are neither constant throughout the course of the application nor similar between subjects (Schoisswohl et al., 2024), other parameters might have contributed to the switch into an excitatory aftereffect.

One of these parameters is the induced current direction with which the stimulation was applied: In the literature standard biphasic pulses with induced AP-PA current direction showed increased mean MEP amplitudes after 1 Hz rTMS in 6 out of 13 participants, whereas only 4 showed an inhibitory response (Goetz et al., 2016). Others stated that 3 out of 4 subjects had inhibitory responses, whereby the induced current direction is not stated (Wassermann et al., 1996). In our experiment only 3 subjects had inhibitory responses to 1 Hz rTMS induced with AP-PA current direction and only one subject to PA-AP. In the literature, monophasic pulses, albeit rectangular, induced higher inhibitory aftereffects than biphasic ones and monophasic pulses inducing AP current direction elicited higher inhibitory aftereffects than pulses with PA current direction (Goetz et al., 2016). Since many conventional TMS pulse sources do not have the capacity to administrate longer lasting monophasic rTMS sessions, this factor should be included in the technical requirements for updates of existing and the development of novel pulse sources. These results indicate that our setup with both biphasic rTMS and probing pulses might suffer from higher variability due to inducing current flows with two directionalities that activate different parts and sets of neurons (Di Lazzaro et al., 2001, 2018) and may thus evoke net stimulation of different cortical regions (Mills et al., 1992). Therefore, the comparability between the two aftereffect measures may have been biased by the different probing as well.

Looking at test-retest reliability per current direction, ICC for induced AP-PA 1 Hz rTMS aftereffect, which was overall no change, was poor with 0.342 and the ICC for the induced PA-AP 1 Hz rTMS aftereffect, which showed excitatory reactions, was moderate with a value of 0.664 (Koo & Li, 2016). Overall, 1 Hz rTMS showed moderate reliability (ICC = 0.561). The range of these reliability values is coherent with previous findings of rTMS reliability and even better within the 1 Hz rTMS comparison of r = 0.266 (Maeda et al., 2000) and r = 0.162 (Prei et al., 2023), which both were only applied with AP-PA current direction. Differences in reliability between the current directions in our results thus may have resulted from the stability of the excitatory effect in the PA-AP condition. Only 9 % of the variability within our results was explained by the best fitting model, as indicated by the conditional *R*^2^ (Nakagawa et al., 2016). Since the explained variability by the fixed effects (marginal *R*^2^ ) reached only 2 %, the individual variability explained circa 7 % of the overall variation of the data (Nakagawa et al., 2016).

The overall cortico-spinal excitability in our study was higher on day one of stimulation with each current direction than on day two, which was exactly one week later. Seven days between sessions should have eliminated cumulative plasticity (Bäumer et al., 2003). Not only did this difference between days in our study contribute to the poor to moderate test-retest reliability of the 1 Hz rTMS aftereffect, but it raises the question on whether participants experienced a habituation effect. Previous literature with an induced current direction of AP-PA reported no differences in aftereffects between days in the 1 Hz rTMS protocol (Maeda et al., 2000) or the tendency to more individual excitation effects for 1 Hz rTMS as depicted in the contingency tables (Prei et al., 2023).

With all these varying outcomes in and possible influences on 1 Hz rTMS (Ridding & Ziemann, 2010), we were interested which parameters from our setup and sample might have contributed to aftereffect differences. With an ordinal logistic regression model, we were able to show that the PA-AP current direction increased the probability to exhibit a more excitatory response. Yet, no other parameters (participants’ sex, attention, excitement and tiredness) reached significance. This finding highlights that although variation in TMS and rTMS responses are well documented (Goldsworthy et al., 2021), influencing factors on both aftereffects and reliability have not been studied thoroughly and need further evaluation, especially with larger sample sizes.

## Conclusions

Contrary to presumes, 2000 pulses of 1 Hz rTMS induced excitatory aftereffects on group level when applied with a current direction of PA-AP in the brain over the healthy motor cortex. When applied with AP-PA current direction, no change in MEP amplitude was detectable. Only three out of thirteen subjects responded to 1 Hz rTMS with cortico-spinal inhibition. Current direction was the only factor to influence the 1 Hz rTMS aftereffect in an ordinal regression model and reliability was poor for AP-PA and moderate for PA-AP current direction. These findings highlight the influence of current direction on 1 Hz rTMS aftereffects and reliability and additionally emphasize that the heuristic regarding the inhibitory effects of low-frequency rTMS should not be applied blindly.

## CRediT authorship contribution statement

Carolina Kanig: Writing – review & editing, Writing – original draft, Methodology, Investigation, Formal analysis, Visualization, Data curation. Mirja Osnabruegge: Writing – review & editing, Methodology, Investigation. Leo Tomasevic: Writing – review & editing, Methodology. Berthold Langguth: Writing – review & editing, Resources. Wolfgang Mack: Writing – review & editing, Supervision, Resources, Funding acquisition. Stefan Schoisswohl: Writing – review & editing, Writing – original draft, Supervision, Funding acquisition, Methodology, Resources, Conceptualization.

## Funding

This work was funded by the dtec.bw – Digitalization and Technology Research Center of the Bundeswehr (MEXT project). dtec.bw is funded by the European Union – NextGenerationEU.

## Declaration of competing interest

The authors declare that they have no known competing financial interests or personal relationships that could have influenced the work reported in this paper.

## Data availability

Data will be made available on request.

## Notes

### Competing Interest Statement

The authors have declared no competing interest.

## References

Agboada, D., Osnabruegge, M., Rethwilm, R., Kanig, C., Schwitzgebel, F., Mack, W., Schecklmann, M., Seiberl, W., & Schoisswohl, S. (2023). Semi-automated motor hotspot search (SAMHS): A framework toward an optimised approach for motor hotspot identification. Frontiers in Human Neuroscience, 17, 1228859. 10.3389/fnhum.2023.1228859

Bates, D., Mächler, M., Bolker, B., & Walker, S. (2015). Fitting Linear Mixed-Effects Models Using lme4. Journal of Statistical Software, 67(1), 1–48. 10.18637/jss.v067.i01

Bäumer, T., Lange, R., Liepert, J., Weiller, C., Siebner, H. R., Rothwell, J. C., & Münchau, A. (2003). Repeated premotor rTMS leads to cumulative plastic changes of motor cortex excitability in humans. NeuroImage, 20(1), 550–560. 10.1016/S1053-8119(03)00310-0

Bech, P., Rasmussen, N. A., Olsen, L. R., Noerholm, V., & Abildgaard, W. (2001). The sensitivity and specificity of the Major Depression Inventory, using the Present State Examination as the index of diagnostic validity. Journal of Affective Disorders, 66(2–3), 159–164. 10.1016/s0165-0327(00)00309-8

Casula, E. P., Rocchi, L., Hannah, R., & Rothwell, J. C. (2018). Effects of pulse width, waveform and current direction in the cortex: A combined cTMS-EEG study. Brain Stimulation, 11(5), 1063–1070. 10.1016/j.brs.2018.04.015

Chen, R., Classen, J., Gerloff, C., Celnik, P., Wassermann, E. M., Hallett, M., & Cohen, L. G. (1997). Depression of motor cortex excitability by low-frequency transcranial magnetic stimulation. Neurology, 48(5), 1398–1403. 10.1212/wnl.48.5.1398

Christensen, R. H. B. (2023). ordinal—Regression Models for Ordinal Data [Computer software]. CRAN. https://CRAN.R-project.org/package=ordinal

Daskalakis, Z. J., Möller, B., Christensen, B. K., Fitzgerald, P. B., Gunraj, C., & Chen, R. (2006). The effects of repetitive transcranial magnetic stimulation on cortical inhibition in healthy human subjects. Experimental Brain Research, 174(3), 403–412. 10.1007/s00221-006-0472-0

Davila-Pérez, P., Jannati, A., Fried, P. J., Cudeiro Mazaira, J., & Pascual-Leone, A. (2018). The Effects of Waveform and Current Direction on the Efficacy and Test-Retest Reliability of Transcranial Magnetic Stimulation. Neuroscience, 393, 97–109. 10.1016/j.neuroscience.2018.09.044

Di Lazzaro, V., Oliviero, A., Pilato, F., Saturno, E., Dileone, M., Mazzone, P., Insola, A., Tonali, P. A., & Rothwell, J. C. (2004). The physiological basis of transcranial motor cortex stimulation in conscious humans. Clinical Neurophysiologyr!: Official Journal of the International Federation of Clinical Neurophysiology, 115(2), 255–266. 10.1016/j.clinph.2003.10.009

Di Lazzaro, V., Oliviero, A., Saturno, E., Pilato, F., Insola, A., Mazzone, P., Profice, P., Tonali, P., & Rothwell, J. C. (2001). The effect on corticospinal volleys of reversing the direction of current induced in the motor cortex by transcranial magnetic stimulation. Experimental Brain Research, 138(2), 268–273. 10.1007/s002210100722

Di Lazzaro, V., Pilato, F., Dileone, M., Profice, P., Oliviero, A., Mazzone, P., Insola, A., Ranieri, F., Tonali, P. A., & Rothwell, J. C. (2008). Low-frequency repetitive transcranial magnetic stimulation suppresses specific excitatory circuits in the human motor cortex. The Journal of Physiology, 586(18), 4481–4487. 10.1113/jphysiol.2008.159558

Di Lazzaro, V., Rothwell, J., & Capogna, M. (2018). Noninvasive Stimulation of the Human Brain: Activation of Multiple Cortical Circuits. The Neuroscientistr!: A Review Journal Bringing Neurobiology, Neurology and Psychiatry, 24(3), 246–260. 10.1177/1073858417717660

Fitzgerald, P. (2002). Intensity-dependent effects of 1 Hz rTMS on human corticospinal excitability. Clinical Neurophysiology, 113(7), 1136–1141. 10.1016/S1388-2457(02)00145-1

Fox, J., & Weisberg, S. (2019). An R companion to applied regression (Third). SAGE Publications. https://www.john-fox.ca/Companion/

Gamer, M., Lemon, J., & Puspendra Singh, I. F. (2019). irr: Various Coefficients of Interrater Reliability and Agreement [Computer software]. CRAN. https://cran.r-project.org/package=irr

Goetz, S. M., Luber, B., Lisanby, S. H., Murphy, D. L. K., Kozyrkov, I. C., Grill, W. M., & Peterchev, A. V. (2016). Enhancement of Neuromodulation with Novel Pulse Shapes Generated by Controllable Pulse Parameter Transcranial Magnetic Stimulation. Brain Stimulation, 9(1), 39–47. 10.1016/j.brs.2015.08.013

Goldsworthy, M. R., Hordacre, B., Rothwell, J. C., & Ridding, M. C. (2021). Effects of rTMS on the brain: Is there value in variability? Cortex; a Journal Devoted to the Study of the Nervous System and Behavior, 139, 43–59. 10.1016/j.cortex.2021.02.024

Gómez-Rubio, V. (2017). ggplot2—Elegant Graphics for Data Analysis (2nd Edition). Journal of Statistical Software, 77(Book Review 2). 10.18637/jss.v077.b02

Guerra, A., López-Alonso, V., Cheeran, B., & Suppa, A. (2020). Variability in non-invasive brain stimulation studies: Reasons and results. Neuroscience Letters, 719, 133330. 10.1016/j.neulet.2017.12.058

Hagströmer, M., Oja, P., & Sjöström, M. (2006). The International Physical Activity Questionnaire (IPAQ): A study of concurrent and construct validity. Public Health Nutrition, 9(6), 755–762. 10.1079/phn2005898

Hastie, T. J. (2017). Statistical Models in S (T. J. Hastie, Ed.; First edition). CRC Press.

Kammer, T., Beck, S., Thielscher, A., Laubis-Herrmann, U., & Topka, H. (2001). Motor thresholds in humans: A transcranial magnetic stimulation study comparing different pulse waveforms, current directions and stimulator types. Clinical Neurophysiology, 112(2), 250–258. 10.1016/S1388-2457(00)00513-7

Kanig, C., Osnabruegge, M., Schwitzgebel, F., Litschel, K., Seiberl, W., Mack, W., Schoisswohl, S., & Schecklmann, M. (2023). Retest reliability of repetitive transcranial magnetic stimulation over the healthy human motor cortex: A systematic review and meta-analysis. Frontiers in Human Neuroscience, 17, 1237713. 10.3389/fnhum.2023.1237713

Kanig, C., Osnabruegge, M., Schwitzgebel, F., Mack, W., Schecklmann, M., & Schoisswohl, S. (2025). Influences of current direction on 1 Hz motor cortex rTMS. Brain Research Bulletin, 230, 111484. 10.1016/j.brainresbull.2025.111484

Klomjai, W., Katz, R., & Lackmy-Vallée, A. (2015). Basic principles of transcranial magnetic stimulation (TMS) and repetitive TMS (rTMS). Annals of Physical and Rehabilitation Medicine, 58(4), 208–213. 10.1016/j.rehab.2015.05.005

Koo, T. K., & Li, M. Y. (2016). A Guideline of Selecting and Reporting Intraclass Correlation Coefficients for Reliability Research. Journal of Chiropractic Medicine, 15(2), 155–163. 10.1016/j.jcm.2016.02.012

Kuhn, M. (2008). Building Predictive Models in R Using the caret Package. Journal of Statistical Software, 28(5), 1–26. 10.18637/jss.v028.i05

Kuznetsova, A., Brockhoff, P. B., & Christensen, R. H. B. (2017). lmerTest Package: Tests in Linear Mixed Effects Models. Journal of Statistical Software, 82(13). 10.18637/jss.v082.i13

Lang, N., Harms, J., Weyh, T., Lemon, R. N., Paulus, W., Rothwell, J. C., & Siebner, H. R. (2006). Stimulus intensity and coil characteristics influence the efficacy of rTMS to suppress cortical excitability. Clinical Neurophysiologyr!: Official Journal of the International Federation of Clinical Neurophysiology, 117(10), 2292–2301. 10.1016/j.clinph.2006.05.030

Lefaucheur, J.-P., Aleman, A., Baeken, C., Benninger, D. H., Brunelin, J., Di Lazzaro, V., Filipović, S. R., Grefkes, C., Hasan, A., Hummel, F. C., Jääskeläinen, S. K., Langguth, B., Leocani, L., Londero, A., Nardone, R., Nguyen, J.-P., Nyffeler, T., Oliveira-Maia, A. J., Oliviero, A., … Ziemann, U. (2020). Evidence-based guidelines on the therapeutic use of repetitive transcranial magnetic stimulation (rTMS): An update (2014–2018). Clinical Neurophysiology, 131(2), 474–528. 10.1016/j.clinph.2019.11.002

Lehrl, S. (2018). Manual zum MWT-B: [Mehrfachwahl-Wortschatz-Intelligenztest] (6. Auflage). Spitta GmbH.

Lüdecke, D., Ben-Shachar, M. S., Patil, I., Waggoner, P., & Makowski, D. (2021). performance: An R Package for Assessment, Comparison and Testing of Statistical Models. Center for Open Science. 10.31234/osf.io/vtq8f

Maeda, F., Keenan, J. P., Tormos, J. M., Topka, H., & Pascual-Leone, A. (2000). Modulation of corticospinal excitability by repetitive transcranial magnetic stimulation. Clinical Neurophysiology, 111(5), 800–805. 10.1016/S1388-2457(99)00323-5

Martin Bland, J., & Altman, DouglasG. (1986). STATISTICAL METHODS FOR ASSESSING AGREEMENT BETWEEN TWO METHODS OF CLINICAL MEASUREMENT. The Lancet, 327(8476), 307–310. 10.1016/S0140-6736(86)90837-8

Mills, K. R., Boniface, S. J., & Schubert, M. (1992). Magnetic brain stimulation with a double coil: The importance of coil orientation. Electroencephalography and Clinical Neurophysiology, 85(1), 17–21. 10.1016/0168-5597(92)90096-T

Modugno, N., Currà, A., Conte, A., Inghilleri, M., Fofi, L., Agostino, R., Manfredi, M., & Berardelli, A. (2003). Depressed intracortical inhibition after long trains of subthreshold repetitive magnetic stimuli at low frequency. Clinical Neurophysiology, 114(12), 2416–2422. 10.1016/S1388-2457(03)00262-1

Nakagawa, S., Johnson, P. C. D., & Schielzeth, H. (2016). The coefficient of determination R2 and intra-class correlation coefficient from generalized linear mixed-effects models revisited and expanded. 10.1101/095851

Nakagawa, S., Johnson, P. C. D., & Schielzeth, H. (2017). The coefficient of determination R2 and intra-class correlation coefficient from generalized linear mixed-effects models revisited and expanded. Journal of the Royal Society, Interface, 14(134). 10.1098/rsif.2017.0213

Oldfield, R. C. (1971). The assessment and analysis of handedness: The Edinburgh inventory. Neuropsychologia, 9(1), 97–113. 10.1016/0028-3932(71)90067-4

Osnabruegge, M., Schwitzgebel, F., Kanig, C., Franke, H., Kerkel, K., Reissmann, A., Langguth, B., Mack, W., Schecklmann, M., & Schoisswohl, S. (2026). Shape of the Pulse: Pulse Width and Current Direction Effects on Motor Evoked Potentials Using a Cobot-assisted Controllable Transcranial Magnetic Stimulation Device. Neuromodulationr⍰: Journal of the International Neuromodulation Society, 29(2), 289–297. 10.1016/j.neurom.2025.08.418

Pascual-Leone, A., Valls-Sole, J., Wassermann, E. M., & Hallett, M. (1994). Responses to rapid-rate transcranial magnetic stimulation of the human motor cortex.

Patel, R., Silla, F., Pierce, S., Theule, J., & Girard, T. A. (2020). Cognitive functioning before and after repetitive transcranial magnetic stimulation (rTMS): A quantitative meta-analysis in healthy adults. Neuropsychologia, 141, 107395. 10.1016/j.neuropsychologia.2020.107395

Peinemann, A., Reimer, B., Löer, C., Quartarone, A., Münchau, A., Conrad, B., & Siebner, H. R. (2004). Long-lasting increase in corticospinal excitability after 1800 pulses of subthreshold 5 Hz repetitive TMS to the primary motor cortex. Clinical Neurophysiologyr⍰: Official Journal of the International Federation of Clinical Neurophysiology, 115(7), 1519–1526. 10.1016/j.clinph.2004.02.005

Pitcher, J. B., Ogston, K. M., & Miles, T. S. (2003). Age and sex differences in human motor cortex input-output characteristics. The Journal of Physiology, 546(Pt 2), 605–613. 10.1113/jphysiol.2002.029454

Prei, K., Kanig, C., Osnabruegge, M., Langguth, B., Mack, W., Abdelnaim, M., Schecklmann, M., & Schoisswohl, S. (2023). Limited evidence for reliability of low and high frequency rTMS over the motor cortex. Brain Research, 1820, 148534. 10.1016/j.brainres.2023.148534

Ridding, M. C., & Ziemann, U. (2010). Determinants of the induction of cortical plasticity by non-invasive brain stimulation in healthy subjects. The Journal of Physiology, 588(Pt 13), 2291–2304. 10.1113/jphysiol.2010.190314

Romero, J. R., Anschel, D., Sparing, R., Gangitano, M., & Pascual-Leone, A. (2002). Subthreshold low frequency repetitive transcranial magnetic stimulation selectively decreases facilitation in the motor cortex. Clinical Neurophysiology, 113(1), 101–107. 10.1016/s1388-2457(01)00693-9

Rossini, P. M., Burke, D., Chen, R., Cohen, L. G., Daskalakis, Z., Di Iorio, R., Di Lazzaro, V., Ferreri, F., Fitzgerald, P. B., George, M. S., Hallett, M., Lefaucheur, J. P., Langguth, B., Matsumoto, H., Miniussi, C., Nitsche, M. A., Pascual-Leone, A., Paulus, W., Rossi, S., … Ziemann, U. (2015). Non-invasive electrical and magnetic stimulation of the brain, spinal cord, roots and peripheral nerves: Basic principles and procedures for routine clinical and research application. An updated report from an I.F.C.N. Committee. Clinical Neurophysiologyr⍰: Official Journal of the International Federation of Clinical Neurophysiology, 126(6), 1071–1107. 10.1016/j.clinph.2015.02.001

Russell V. Lenth. (2023). emmeans: Estimated Marginal Means, aka Least-Squares Means [Computer software]. https://CRAN.R-project.org/package=emmeans

Russo, S., Sarasso, S., Puglisi, G. E., Dal Palù, D., Pigorini, A., Casarotto, S., D’Ambrosio, S., Astolfi, A., Massimini, M., Rosanova, M., & Fecchio, M. (2022). TAAC -TMS Adaptable Auditory Control: A universal tool to mask TMS clicks. Journal of Neuroscience Methods, 370, 109491. 10.1016/j.jneumeth.2022.109491

Sack, A. T., Paneva, J., Küthe, T., Dijkstra, E., Zwienenberg, L., Arns, M., & Schuhmann, T. (2024). Target Engagement and Brain State Dependence of Transcranial Magnetic Stimulation: Implications for Clinical Practice. Biological Psychiatry, 95(6), 536–544. 10.1016/j.biopsych.2023.09.011

Schoisswohl, S., Kanig, C., Osnabruegge, M., Agboada, D., Langguth, B., Rethwilm, R., Hebel, T., Abdelnaim, M. A., Mack, W., Seiberl, W., Kuder, M., & Schecklmann, M. (2024). Monitoring Changes in TMS-Evoked EEG and EMG Activity During 1 Hz rTMS of the Healthy Motor Cortex. eNeuro, 11(4). 10.1523/ENEURO.0309-23.2024

Schoisswohl, S., Langguth, B., Hebel, T., Abdelnaim, M. A., Volberg, G., & Schecklmann, M. (2021). Heading for Personalized rTMS in Tinnitus: Reliability of Individualized Stimulation Protocols in Behavioral and Electrophysiological Responses. Journal of Personalized Medicine, 11(6), 536. 10.3390/jpm11060536

Schoisswohl, S., Langguth, B., Weber, F. C., Abdelnaim, M. A., Hebel, T., Mack, W., & Schecklmann, M. (2023). One way or another: Treatment effects of 1 Hz rTMS using different current directions in a small sample of tinnitus patients. Neuroscience Letters, 797, 137026. 10.1016/j.neulet.2022.137026

Sommer, M., Norden, C., Schmack, L., Rothkegel, H., Lang, N., & Paulus, W. (2013). Opposite optimal current flow directions for induction of neuroplasticity and excitation threshold in the human motor cortex. Brain Stimulation, 6(3), 363–370. 10.1016/j.brs.2012.07.003

Spampinato, D. A., Ibanez, J., Rocchi, L., & Rothwell, J. (2023). Motor potentials evoked by transcranial magnetic stimulation: Interpreting a simple measure of a complex system. The Journal of Physiology, 601(14), 2827–2851. 10.1113/JP281885

Touge, T., Gerschlager, W., Brown, P., & Rothwell, J. C. (2001). Are the after-effects of low-frequency rTMS on motor cortex excitability due to changes in the efficacy of cortical synapses? Clinical Neurophysiology, 112(11), 2138–2145. 10.1016/s1388-2457(01)00651-4

Wassermann, E. M., Grafman, J., Berry, C., Hollnagel, C., Wild, K., Clark, K., & Hallett, M. (1996). Use and safety of a new repetitive transcranial magnetic stimulator. Electroencephalography and Clinical Neurophysiology/Electromyography and Motor Control, 101(5), 412–417. 10.1016/0924-980X(96)96004-X

Weisz, N., Steidle, L., & Lorenz, I. (2012). Formerly known as inhibitory: Effects of 1-Hz rTMS on auditory cortex are state-dependent. The European Journal of Neuroscience, 36(1), 2077–2087. 10.1111/j.1460-9568.2012.08097.x

